# Analysis of IGHA1 and other salivary proteins post half marathon in female participants

**DOI:** 10.1101/2022.11.13.515896

**Authors:** Yosuke Maruyama, Tomoaki Seki, Seiichi Ando, Hiroki Tanabe, Hitoshi Mori

## Abstract

**Background:** High intensity exercise, such as in marathons and triathlons, suppresses transient local and systemic immunity. Much is known about the systemic immunosuppression response, but little is known about its local response in the oral cavity, lungs, bronchial tubes, and skin. The oral cavity is an entrance for bacteria or viruses to enter the body. Saliva covers the epidermis of the oral cavity and plays an important role in the local stress response by preventing infection. In this study, we examined the properties of saliva secreted during the local stress response induced by a half marathon (HM) using quantitative proteomic analysis.

**Methods:** The Exercise group (ExG), 19 healthy female university students participated in (HM) race. The non-exercise group (NExG), 16 healthy female university students had not participated in the ExG. In ExG, saliva samples were collected 1h pre and 2h and 4h post HM. NExG were collected in same time interval. Saliva volume, protein concentration and relative IGHA1 expression were analyzed. In addition, 1h pre and 2h post HM saliva were analyzed by iTRAQ. The identified factors of iTRAQ, analyzed for the ExG and NExG by western blotting.

**Results:** We identified kallikrein 1 (KLK1), immunoglobulin kappa chain (IgK) and cystatin S (CST4) as suppression factors, as well as immunoglobulin heavy constant alpha 1 (IGHA1), which has been reported as an immunological stress marker. KLK1 (*p* = 0.011), IGK (*p* = 0.002), and CST4 (*p* = 0.003) were suppressed 2 h post-HM compared to pre-HM, and KLK1 (*p* = 0.004) and CST4 (*p* = 0.006) were suppressed 4 h post-HM. There was also a positive correlation between IGHA1, IGK, and CST4 2 and 4 h post-HM. In addition, KLK1 and IGK after 2 h post-HM showed a positive correlation.

**Conclusion:** Our study demonstrated that the salivary proteome is regulated and antimicrobial proteins are suppressed post-HM. These results suggest that oral immunity was transiently suppressed post-HM. The positive correlation of each protein at 2 and 4 h post-HM suggests that the suppressed state was similarly regulated up to 4 h after a HM. The proteins identified in this study may have applications as stress markers for recreational runners and individuals who perform high- and moderate-intensity exercises in their daily lives.

## Introduction

Moderate exercise, such as 30 minutes of walking, daily running, and regular sports, has been reported to reduce the risk of upper respiratory tract infections (URTI) (Peters E. M., 1997; Nieman et al., 2011; Bigley et al., 2013; Antunes et al., 2016). Moderate exercise decreases inflammatory cytokines and oxidative stress and improves the function of various immune cells in the resting state (Peters E. M., 1997; Abd El-Kader et al., 2018). On the other hand, prolonged and transient high-intensity exercise (HIE) and training, such as marathons or triathlons, may suppress the immune system and increase the risk of developing URTI (Nieman et al., 1990; Fitzgerald L., 1991; Heath et al., 1991). The risk of URTI and lower respiratory tract infections after HIE is higher in healthy individuals than in athletes (Cantó et al., 2018), and it has been reported that there is no difference in risk between males and females (Gleeson et al., 2011; Cantó et al., 2018). The open-window theory explains the temporary immunosuppression caused by high-intensity or extensive exercise that leads to exhaustion. The risk of URTI due to immunosuppression of exercise intensity has also been explained using the J-curve model (Nieman, 2000). A comparison of blood component data before and after marathons showed a decline in the quantitative and qualitative function of immune cells responsible for innate immunity, such as lymphocytes, natural killer (NK) cells, and macrophages, as well as a decline in the function of cellular immunity between 3 and 72 h after participation in a marathon (Nieman, D. C. & Wentz, 2019). Regarding blood components during immunosuppression by HIE, it has been reported that there is no difference between males and females in plasma immunoglobulin A (IgA), immunoglobulin G and immunoglobulin M concentrations, total white blood cell count, neutrophil count, monocyte count, and lymphocyte count (Gleeson et al., 1999). On the other hand, it has been found that there are differences between men and women in salivary flow rate, IgA concentration, IgA secretion, B cell count, and NK cell count (Gleeson et al., 2011). Depending on the exercise intensity, immunosuppression after HIE has been observed for 2–24 h after exercise, and recovery to the pre-exercise state occurs for approximately two to three days (Fitzgerald L., 1991; Kakanis et al., 2010). HIE causes both systemic and local immunosuppression (e.g., in the oral cavity, lungs, bronchial tubes, and skin) after exercise. On the other hand, participation in a half marathon (HM) or middle intensity exercise (not sufficient to induce systemic immunosuppression) has been reported to cause muscle injury and systemic and local immunosuppression (Briviba et al., 2005; Lippi et al., 2011).

The oral cavity is the first fronttal region where bacteria and viruses encounter the body and plays an important role in preventing infection. HIE decreases the secretion of saliva and antimicrobial substances (Mackinnon et al., 1993; Nieman et al., 2002). Saliva is secreted by the parotid, submandibular, and sublingual glands and numerous minor salivary glands (Edgar, 1992). When saliva is supplied to the oral cavity, it provides mucin, lactoferrin, IgA, lysozyme, and lactoperoxidase, which have been reported to contribute to antibacterial activity (Amerongen et al., 2002). Salivary volume and secretion of salivary components are controlled by the autonomic nervous system (Proctor G. B. & Carpenter, 2007). Exercise affects saliva secretion by activating the parasympathetic nervous system, thereby regulating the salivary glands. Short-term exercise increases salivary secretory IgA (sIgA), whereas strenuous exercises, such as interval training, marathons, and cycling races, have been reported to decrease salivary flow and sIgA secretion (Mackinnon et al., 1993; Nieman et al., 2002; Klentrou et al., 2002; Laing et al., 2005). IgA is a glycoprotein produced by mature B cells in blood (Macpherson et al., 2008). It is secreted into bodily fluids such as tears and saliva, as well as bronchial, nasopharyngeal, intestinal, and urogenital secretions (Klentrou et al., 2002). IgA is composed of two heavy chains and two light chains. The immunoglobulin heavy constant alpha 1 (IGHA1) gene encodes a constant region of immunoglobulin heavy chain. The immunoglobulin kappa light chain (IGK) and immunoglobulin lambda light chain (IGλ) genes encode a constant region of immunoglobulin light chains (Woof et al., 2011). IgA can form dimers, which are stabilized between the heavy chains of each monomer and joining site and the J chain (Woof et al., 2011). Salivary IgA is used as a mental and physical stress marker (Pedersen et al., 2019). The reported roles of IgA include the inhibition of bacterial aggregation, neutralization of viruses, and opsonization of support (Kaetzel et al., 1991; Walker D. M., 2004; Fábián et al., 2012).

Decreased sIgA secretion due to high training load is thought to be associated with an increased risk of respiratory disease (Fahlman et al., 2005; Neville et al., 2008; Walsh et al., 2011; Leicht et al., 2012). Previous studies on the immunosuppressive state induced by exercise have mainly used the proteome and metabolome of blood components. Consequently, the detailed behavior of blood metabolites has become clearer (Kurgan et al., 2019; Guseh et al., 2020). However, studies on the oral cavity have been confined to quantifying salivary volume, protein concentration, and marker proteins to easily determine the body’s immune status. Few studies have quantitatively analyzed salivary proteins before and after the immune suppression state in the oral cavity, which is the frontline of defense response to pathogen infection.

In this study, we performed a quantitative proteomic analysis of saliva in the immunosuppressed state (i.e., decreased salivary outflow, increased salivary protein concentration, and decreased IgA per unit protein) in the oral cavity induced by a HM and tested the correlation of regulated proteins. The results provide insight into the local stress state induced by a HM. Furthermore, this insight into local stress responses may contribute to the health management of athletes and amateur recreational runners temporarily exposed to exercise stress.

## Materials & Methods

### Participants and experimental design

In the exercise group (ExG), 19 healthy female university students participated in HM race held in Shibetsu City, Hokkaido (44°10’ N, 144°24’ E) in 2018. The participants were volunteers with no known underlying diseases and predisposing factor to infection. They trained for approximately two months prior to the HM, including running about 5 – 10 km a week, for endure the HM race. The median and whole range age, weight, body mass index (BMI), HM completion time, Race pace and energy consumption during pre HM to post 2 h HM of the participants were shown in Table 1. In the non-exercise group (NExG), 16 healthy female university students had not participated in the ExG. The sampling of the NExG was performed during the time they attended lectures at the university (Fig. 1). The median and whole range age, weight, BMI and energy consumption during 1^st^ to 2^nd^ sampling were shown in Table 1. The ethics committee of the Nayoro City University approved the experimental protocol (approval number 17-22). All participants gave signed informed consent.

**Table 1.** Participants characteristics. HM energy consumption represented energy consumption during HM in ExG. Energy consumption represented total amount of pre-HM to 2 h post-HM in ExG and 1^st^ to 2^nd^ sampling in NExG. Data show means and standard deviation. The *p*-values refer to the comparison between the NExG and ExG using Mann-Whitney rank test. BMI, Body Mass Index

|  | Median (min - max) |  |  |
| --- | --- | --- | --- |
|  | Ex group (ExG) N=19 | NEx group (NExG) N=16 | *p* value |
| Age (years) | 20.4 (19.1 - 21.4) | 20.1 (18.1 - 22.5) | 0.960 |
| Body mass (kg) | 53.1(44.8 - 67.0) | 49.5 (44.2 - 68.2) | 0.689 |
| Length (cm) | 159 (151 - 177) | 156 (147 - 178) | 0.430 |
| BMI (kg / m<sup>2</sup>) | 20.96 (19.65 - 22.34) | 20.52 (19.65 - 22.34) | 0.401 |
| Running time (min) | 135.05 (107.20 - 148.78) | - | - |
| Race pace (m / min) | 159.8 (142.50 - 188.30 ) | - | - |
| HM Energy consumption (Kcal) | 1152.5 (969.4 - 1583.0) | - | - |
| Average HM Energy consumption (Kcal / min) | 8.58 (6.97-12.72) | - | - |
| Total Energy consumption (Kcal) | 1705.12 (1445.30 - 2419.98) | 588.55 (537.87 - 793.83) | < 0.001 |

**Fig. 1.**
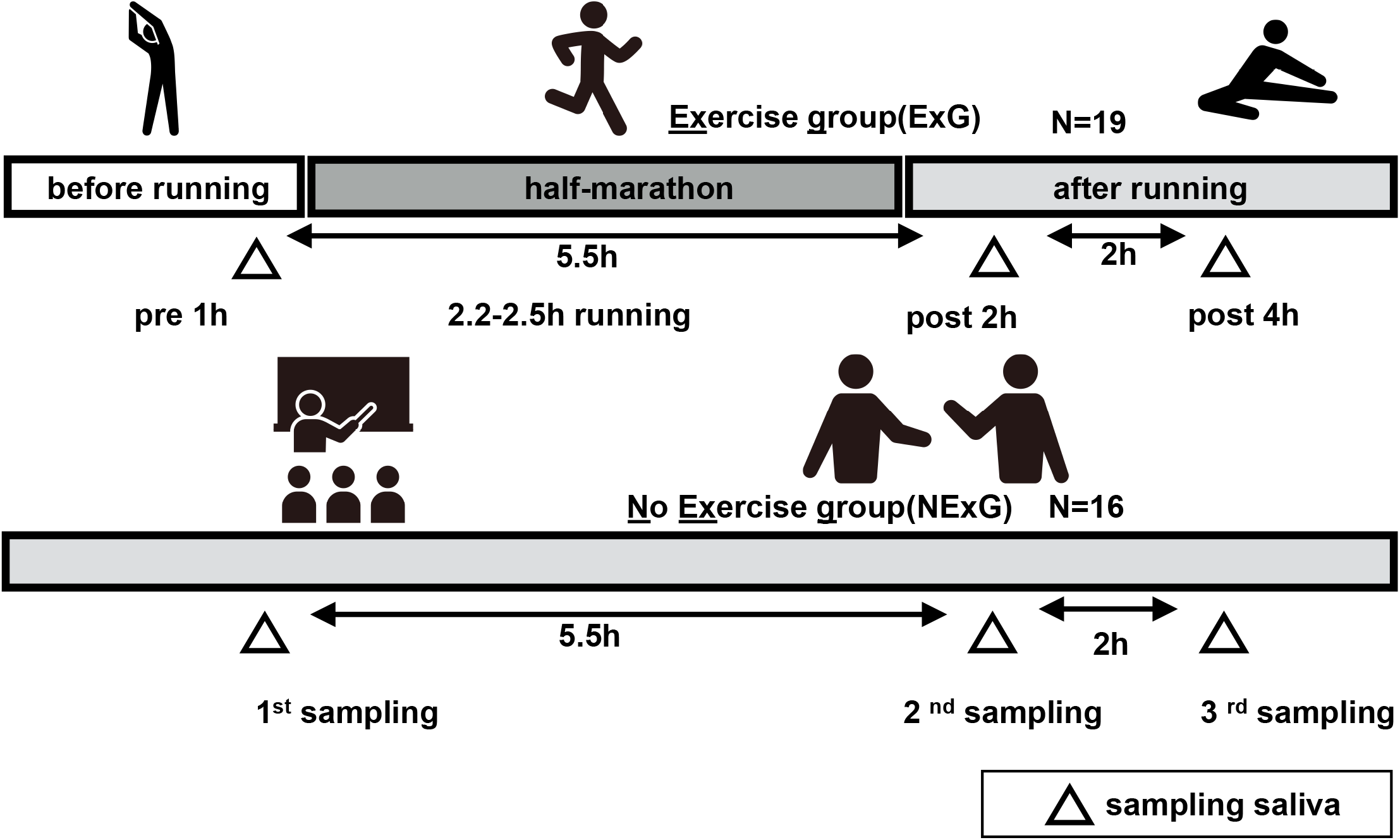
Study diagrams and sampling points of saliva. The exercise group (ExG) that performed the half marathon (HM) and the non-exercise group (NExG) consisted of 19 and 16 participants, respectively. Salivary samples in the ExG were collected 1 h pre HM and post 2 h and post 4h HM, while those in the NExG were also collected at three intervals, 5.5 h and 7.5 h after the first sampling. The intervals between the effect of the HM on secretion rate, protein concentration, and IgA secretion in saliva were approximately the same.

### Saliva sampling

Salivette Cotton (Sarstedt K. K, 51.1534, Sarstedt, Germany) was used for saliva sampling. Saliva was collected from the participants that rinsed their mouths for 5 min with water several times before sampling. At the sampling time, the patients put in mouth a cotton roll for the duration of 2 minutes. The ExG was sampled at 1 h pre, 2 h post, and 4 h post-HM. For the NExG, the first sampling was performed at 10:00, the second at 14:30, and the third at 18:30 (Fig. 1). The intervals between salivary collections in the ExG and NExG were approximately the same. Water was provided ad libitum in both groups. Saliva samples were fixed in liquid nitrogen and stored at – 80 °C until analysis.

### Analysis of saliva volume and concentration

Saliva volume was calculated by weighing the pre collection tube and post collection tube assuming a density of 1 g/ mL. Saliva flow rates were determined by dividing the volume of saliva by the collection time. Protein concentration was measured by Bradford quantitation method (Bradford, M. M. 1976). We used the Bradford Protein Assay Kit (TAKARA, T9310A, Shiga, Japan).

### SDS-PAGE, Silver Staining and CBB staining

The saliva protein was suspended in 2x SDS sample buffer (200 mM Tris-HCl pH 6.8, 400 mM DTT, 8% SDS, 0.4% bromophenol blue, 40% glycerol), and swept on the 10 – 12.5 % acrylamide gel and separated by sodium dodecyl sulfate-polyacrylamide gel electrophoresis (SDS-PAGE) under reducing conditions. The silver staining was then performed using a silver staining kit (WAKO 291-5030) according to the manufacturer’s protocol. The coomassie brilliant blue (CBB) staining were used as a loading control. CBB staining performed using a EzStain Aqua (ATTO, AE-1340, Tokyo, Japan) according to the manufacturer’s protocol.

### Protein preparation and iTRAQ labeling

Salivary proteins from nine ExG participants were used for the iTRAQ analysis. The iTRAQ technology for protein quantitation using mass spectrometry is a recent, means of determining relative protein levels in up to 4-8 samples simultaneously (Ross et al 2004). A total of 5 μg of each protein was subjected to trypsin treatment using a captured trypsin column (TAKARA, 635722, Shiga, Japan). After trypsin treatment, the samples were run on a mono-spin C18 column (GLS 5010-21732, Tokyo, Japan). The digested peptides were labeled with iTRAQ reagent according to the manufacturer’s protocol (Applied Biosystems). In brief, the digested peptides were mixed with iTRAQ reagents (AB Sciex, 4352135, CA, USA) as follows: 1 h pre-HM (tag 114 or 116) and 2 h post HM (tag 115 or 117) (Fig. 4). The mixed samples were incubated at 24□ for 1 h, and then the pooled samples 114, 115, 116, and 117 were further pooled into one tube. Each sample was treated with an SCX column (GLS, 5010-21726, Tokyo, Japan) and eluted with a cation exchange buffer (10 mM KH_2_PO_4_, 25% acetonitrile). The eluted solution containing the peptides was dried using a vacuum concentrator. The precipitate was dissolved in a solution (2% acetonitrile and 0.1% formic acid) and loaded onto an Eksigent microLC 200 (Eksigent, Dublin, USA) equipped with an electrospray ionization triple quadrupole-linear ion trap system (Triple Quad 5500+LC-MS/MS system, SCIEX). iTRAQ data were analyzed using ProteinPlot™ software ver 5.0.2 (SCIEX). The SwissProt database containing 20411 human sequences, was used for protein verification.

### Western blotting

Salivary samples were centrifuged for 5 min at 12000g at 4 °C before SDS-PAGE. Western blotting (WB) was performed as previously described (Lüllau et al 1996). Briefly, 3 μg of salivary protein was loaded onto 10–12.5% SDS acrylamide gels and separated for 80 min at 30 mA constant current. After SDS-PAGE, proteins were transferred to PVDF membranes (Merck Millipore, IEVH00005, Tokyo, Japan), blocked with 2.5% non-fat milk in PBS containing 0.1% Tween 20 for 1 h, and probed with primary antibodies against IGHA1 (Monoclonal rabbit 1:5000 dilution; 31-1030-00, RevMab Biosciences, South San Francisco, USA), Cystatin S (CST4) (polyclonal rabbit1:5000 dilution; GTX100690, GeneTex, USA), IGK (Polyclonal rabbit 1:2000 dilution; 14678-1-AP, Proteintech Group, Inc, Chicago USA), Zinc Alpha 2 Glycoprotein (ZA2G) (polyclonal rabbit 1:5000 dilution; AP6628a abcepta, CA, USA), and Kallikrein 1 (KLK1) (polyclonal rabbit 1:3000 dilution; CSB-PA005798, Cusabio Technology llc, TX, USA). The specific proteins were detected with horseradish peroxidase (HRP) conjugated secondary antibodies (1:5000 dilution; CtNo_NA934, GE Healthcare, 1:5000 dilution; CtNo_NA931, GE Healthcare, Tokyo, Japan), developed with Immobilon Western Chemiluminescent HRP Substrate (CtNo_WBKLS0100, Merck Millipore, Tokyo, Japan), and visualized using LumiVision PRO 400EX system (AISIN, Aichi, Japan).

### Statistical analysis

The data are presented as median with interquartile range or whole range, as appropriate. All data analyzed as not normally distributed data. The age, height, weight, BMI and total energy consumption were evaluated by Mann-Whitney U-tests. The significant differences in salivary volume, protein concentration, iTRAQ, and WB analyses were examined using the Wilcoxon signed-rank sum test. The statistical significance was set at *p* < 0.05. Statistics were performed with SPSS software (SPSS Inc., IBM,). The Spearman’s rank correlation coefficient test was performed to check the correlation of regulated proteins at pre and post-HM. The boxplots were generated using BoxPlotR (Spitzer et al 2014). The Heatmap was plotted by http://www.bioinformatics.com.cn/srplot, an online platform for data analysis and visualization.

## Results

### HM led to decreased salivary secretion and IGHA1

In this study, 19 participants took part in an HM as the ExG, and 16 participants did not take part in the HM and were the NExG (Fig. 1). In the ExG, salivary secretion was reduced and the amount of salivary proteins increased post-HM compared to those observed pre-HM (Fig. 2A and 2B). There were no differences in salivary volume and protein concentration in the NExG for the 2^nd^ and 3^rd^ sampling intervals compared with the 1^st^ sampling saliva (Fig. 2A and 2B). The amounts of salivary IGHA1 from 2 and 4 h post-HM decreased by a median 0.82- and 0.90-fold, respectively, compared with that observed pre-HM (2h *p* = 0.003, 4h *p* < 0.001). There were no differences in the relative amounts of IGHA1 in NExG at the three sampling intervals (Fig. 2D).

**Fig. 2.**
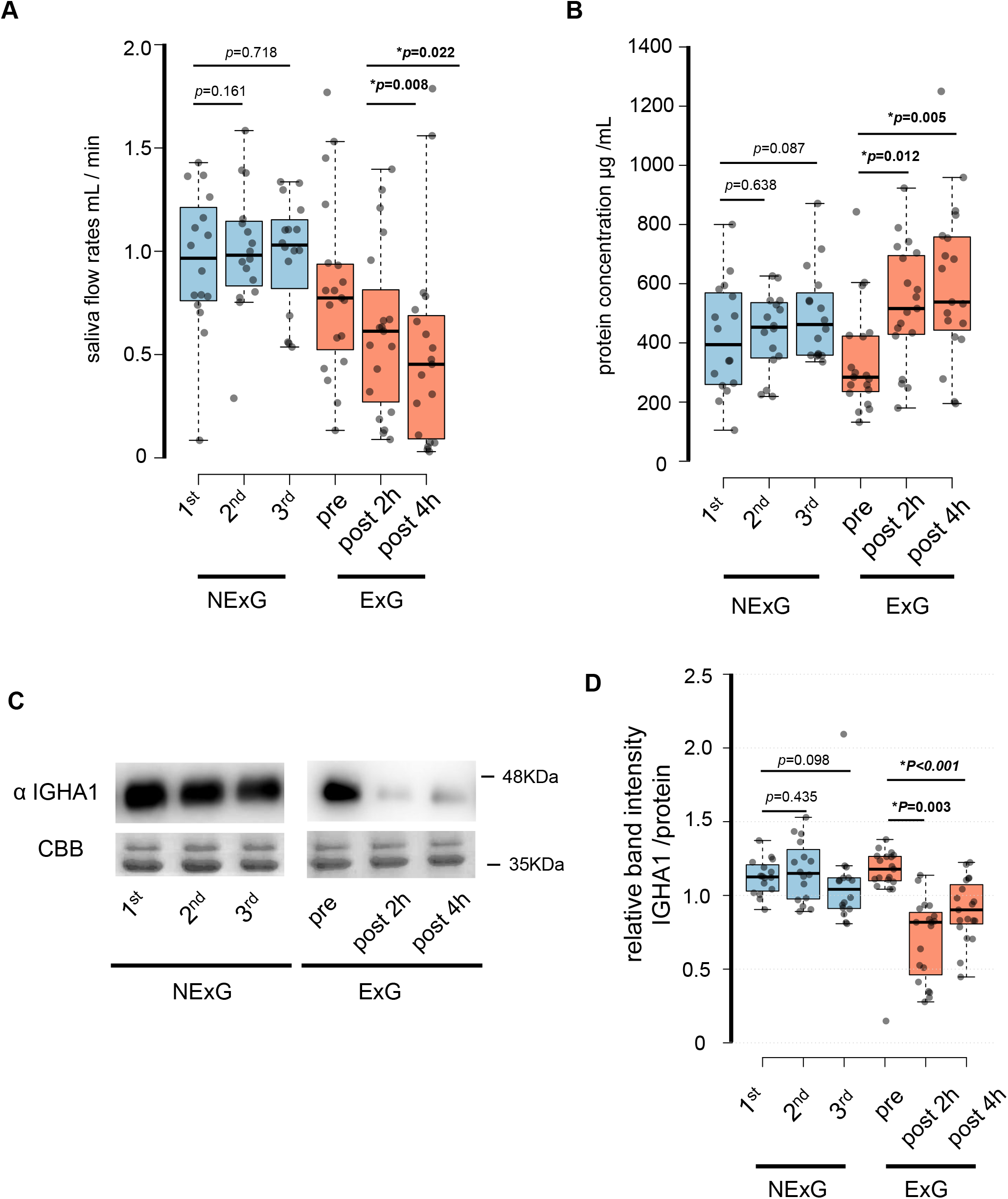
Effect of half-marathon on saliva secretion rate, protein concentration, and IGHA1 secretion. (A) The secretion rate of saliva in the non-exercise group (NExG, N = 16) and exercise group (ExG, N = 19). (B) The salivary protein concentration in the NExG and ExG. (C) Western blot analysis of salivary IGHA1. CBB staining was used as a loading control. (D) The relative band intensity of salivary IGHA1 in the NExG and ExG. The band intensity from first sampling and pre HM was compared with those from second and third samplings in the NExG, and 2 h post and post 4 h HM, respectively. The fold changes were considered as 1.0 at first sampling in the NExG and pre HM in the ExG, respectively. *significant difference at *p* < 0.05 level.

### Proteomic analysis of salivary proteins before and after exercise

To consider the changes in salivary proteins pre- and post-HM in the ExG, we compared salivary proteins separated by SDS-PAGE and performed silver staining (Fig. S1). In the ExG, the protein bands in the vicinity of 15, 30, and 50 kDa decreased in the salivary samples collected 2 and 4 h post-HM compared with pre-HM (Fig. S1). In contrast, no differences in the band intensity of salivary proteins were found at the three sampling intervals of NExG (Fig. S1). To analyze the protein changes pre-and post-HM in the ExG in detail, we performed quantitative proteomic analysis using iTRAQ labeling and examined salivary proteins 1 h pre- and 2 h post-HM in nine participants (Fig. 3A).

**Fig. 3.**
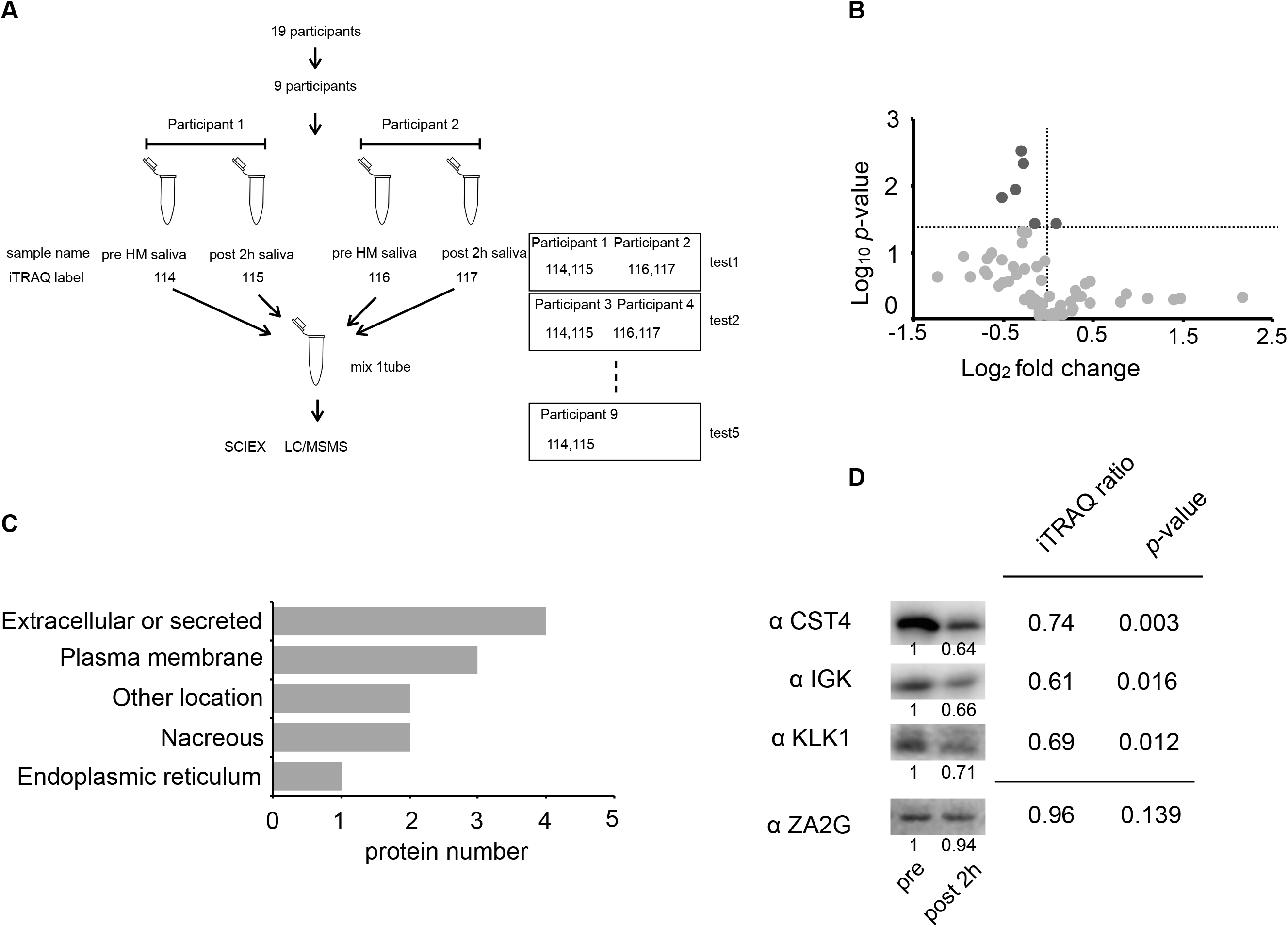
Identification of the half-marathon regulated salivary proteins by iTRAQ analysis. (A) Schematic diagram of the iTRAQ based on experiments. The salivary proteins of 9 participants from pre- and post 2 h HM were analyzed on 5 separate occasions. Four samples were pooled into one sample for either pre-HM and 2 h post-HM. (B) Protein distribution in accordance with log_10_ *P*-versus and log_2_ ratio of the detected 1 ≤ peptide proteins. The darkgray dot indicated the salivary proteins that changed 2 h post-HM compared with pre-HM (*p* < 0.05). (C) Subcellular annotation of differentially expressed proteins in saliva. The subcellular location was shown by UniProt annotation. (D) Western blot profile for CST4, IGK, KLK1, and ZA2G using iTRAQ samples. The numbers below the blot indicate the relative signal intensity of each protein compared with pre-HM sample.

**Fig. 4.**
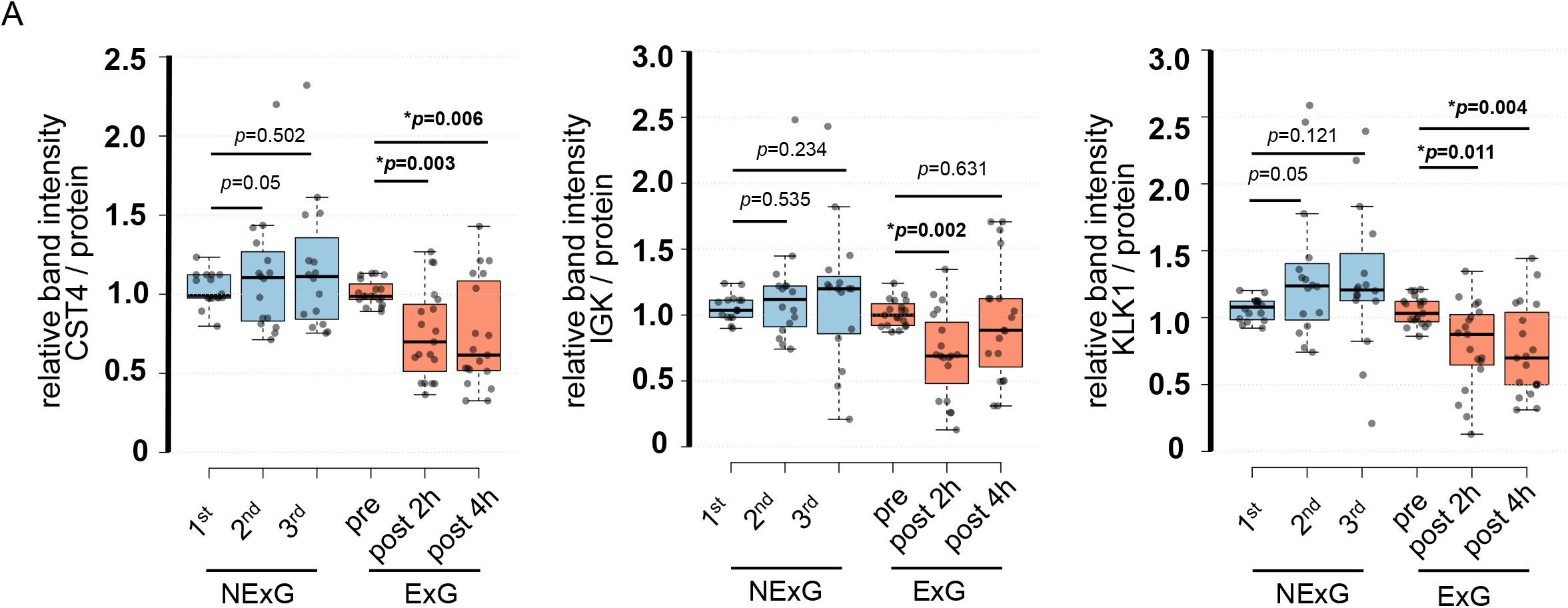
Confirmation of iTRAQ ratio by western blot analyses. (A) The expression of CST4, IGK and KLK1 in saliva. The center lines show the medians of the data in the ExG (N = 19) and NExG (N = 16); box limits indicate the 25^th^ and 75^th^ percentiles as determined by R software; whiskers extended 1.5 times the interquartile range from the 25^th^ and 75^th^ percentiles; outliers are represented by dots. Differences between pre-and post-HM or first and other time points of the NExG were tested using the Wilcoxon signed-rank test. *significant difference at *p* < 0.05 level.

We identified 4089 proteins, as shown in Supplemental Fig. 2, and our iTRAQ data were compared with the protein expression data available through the Human Salivary Protein Wiki (HSP-Wiki: https://salivaryproteome.nidcr.nih.gov/) (Lau W. et al., 2021). The 1039 proteins were matched with the Human Salivary Proteome Wiki data. The 48 proteins were identified as ≤ 1 peptides (Supplemental Table 2 and Fig. 3B), and 6 proteins were significantly different between 1 h pre- and 2 h post-HM (*p* < 0.05) (Fig. 3B). We were also able to identify four proteins that were suppressed less than 0.8-fold, while no proteins that were expressed more than 1.2-fold could be identified (Table 2).

**Table 2.** The salivary proteins regulated by half marathon. The fold change shows the relative ratio of 2 h post-HM to pre-HM. The peptide shows the maximum detected number in analyses samples. *significant difference at *p* < 0.05 level.

| Accession | description | gene name | MW (KDa) | fold change | peptide | *p* value |
| --- | --- | --- | --- | --- | --- | --- |
| P01036 | Cystatin S | *CST4* | 16.21 | 0.74 | 4 | 0.003 |
| P01876 | Immunoglobulin heavy constant alpha 1 | *IGHA1* | 37.65 | 0.76 | 7 | 0.005 |
| P06870 | Kallikrein 1 | *KLK1* | 28.89 | 0.69 | 4 | 0.012 |
| P0DOX7 | Immunoglobulin kappa light chain | *IGK* | 23.37 | 0.60 | 6 | 0.016 |

The factors that were suppressed by less than 0.8-fold were CST4, IGHA1, KLK1, and IGK, with values of 0.74, 0.76, 0.69, and 0.60 post-HM, respectively (Table 2 and Fig. 3B). IGHA1 has been used as a suppression marker of salivary immunity, and similar results were obtained by WB and iTRAQ analyses (Fig. 2C and 2D).

Based on the results of the iTRAQ experiments, we immunoblotted four proteins: CST4, IGK, KLK1, and ZA2G. ZA2G was a control marker with constant expression, whereas the other three proteins were suppressed as variable factors (Fig. 3D).

### Subcellular and functional characterization of the candidate proteins

Among the 48 factors (peptides ≥ 1), 16 were classified as secretory proteins (Table S1). Ten factors (*p* < 0.05) were classified based on UniProt annotation (https://www.uniprot.org/). The four proteins were classified as extracellular or secreted, three as plasma membrane, two as nacreous, and two belonged to other locations (Fig. 3C).

### IGK, CST4, and KLK1 were suppressed by the HM

To further confirm the validation of the iTRAQ results, we examined protein expression levels by WB. WB was performed using saliva collected from the ExG and NExG. The median and *P* values of KLK1 (0.85, *P* = 0.011) and CST4 (0.70, *P* = 0.003) were suppressed 2 h post-HM, respectively, compared with those 1 h pre-HM (Fig. 4). The suppression of KLK1 (0.70, *P* = 0.004) and CST4 (0.62, *P* = 0.006) was observed 4 h post-HM, the same as 2 h post-HM. The IGK (0.69, *P* = 0.002) at 2 h post-HM was suppressed, but that at 4 h post-HM was not (Fig. 4). These results indicate that the iTRAQ ratios were consistent with the quantitative results obtained by WB analysis. In contrast to the ExG, there were no significant differences in these factors at each sampling point of NExG.

### Correlation of IGHA1, IGK, CST4, and KLK1

The relative abundance of protein changes in the ExG was tested using Spearman’s correlation conflict test. A positive correlation was observed between 2 and 4 h post-HM. The *r* and *p* values of IGHA1 (*r* = 0.70, *P* = 0.001), CST4 (*r* = 0.51, *p* = 0.026), and IGK (*r* = 0.46, *p* = 0.049) were observed (Fig. 5). Similarly, a positive correlation was observed between KLK1 protein abundance in 2 h post-HM and IGK 2 h (*r* = 0.72, *p* < 0.001) and IGK 4 h (*r* = 0.64, *p* < 0.001). In contrast, no relationships were observed between other protein abundance and time points (Fig. 5).

**Fig. 5.**
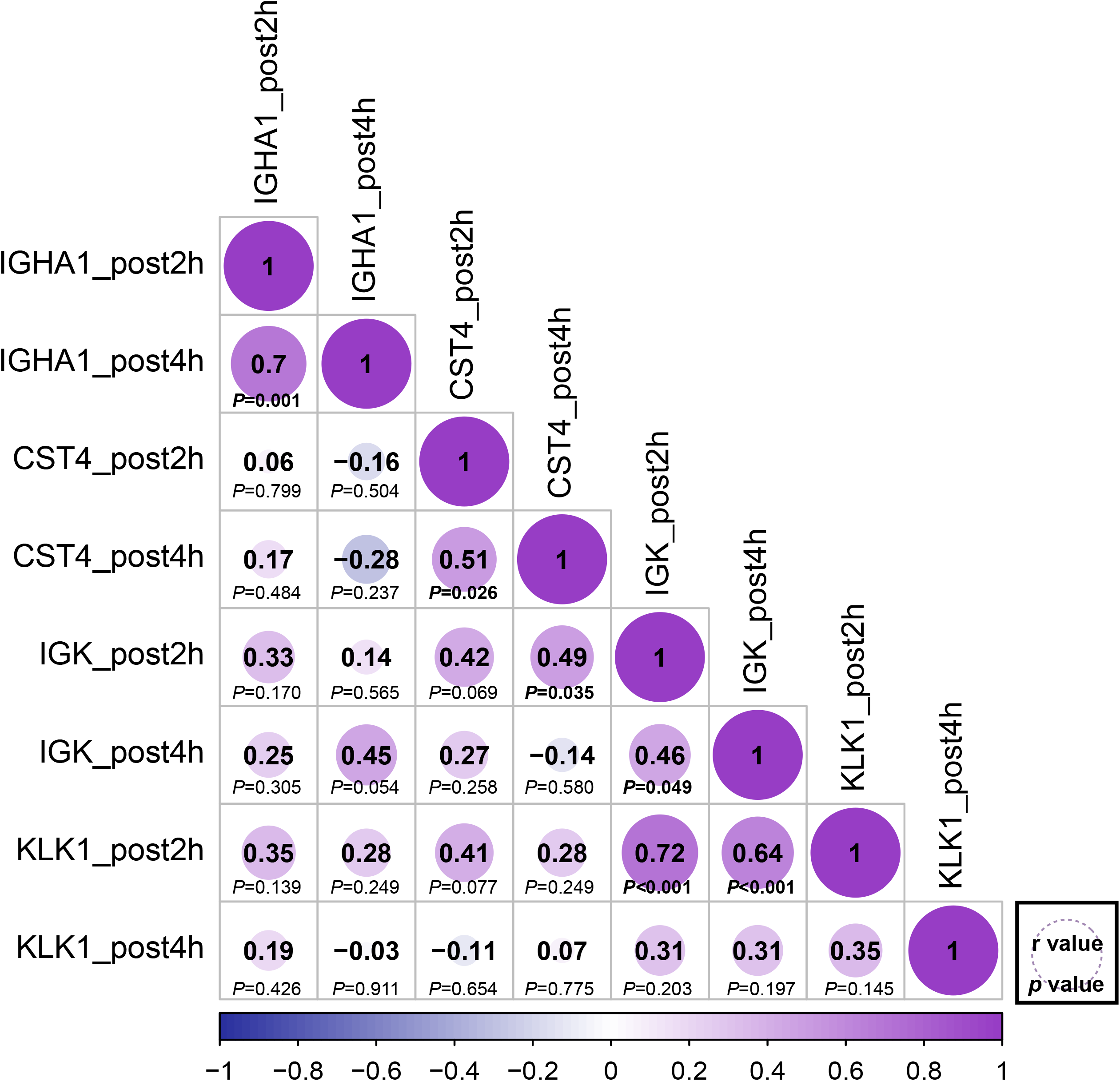
Correlation matrix of different expression protein. Correlations between the relative abundance of IGHA1, CST4, IGK and KLK1 at 2 h post and 4 h post-HM. Spearman correlation coefficients (*r*) were shown for each protein. The bold font *p* - value was shown for significant difference at *p* < 0.05 level.

## Discussion

The findings of this study reveal the salivary proteome during HM stress in the oral cavity. After 2 h, HM stress suppressed salivary antimicrobial proteins such as KLK1, CST4, and IGK in association with IGHA1. After 4 h, HM, IGHA1, KLK1, and CST4, but not IGK, were suppressed after 2 h post-HM.

In addition, the relative abundances of IGHA1, CST4, and IGK were positively correlated at 2 and 4 h post-HM. Similarly, KLK1 after 2 h post-HM was positively correlated with IGK after 2 and 4 h post-HM. The suppression of these antimicrobial proteins suggests that HM suppresses local immunity in the oral cavity and that it lasts from 2 to 4 h post-HM. Our findings will help further the understanding of local stress responses in the oral cavity due to exercise stress.

In the ExG, HM median and whole range running time (135.05, 107.20–148.78 min, respectively,), race pace (132.07, 118.64–139.07 m/min, respectively), and during HM calorie expenditure (1152.5, 969.4–1583.0 kcal, respectively) were shown in Table 1. The running time and race pace were comparable to those reported for women (N = 2852) in the 2017 HM race in Ljubljana, Slovenia (Nikolaidis et al., 2019). We also analyzed the IGHA1 secretion rate, saliva secretion per unit of time, and saliva concentration as immunosuppression markers. The results suggest the induction of the suppression of local immunity in female university students without a general exercise habit. In contrast, there were no changes in the saliva flow rate, concentration, and IGHA1 amount in the normal state of university students taking classes (Fig. 2). Quantitative proteomic analysis using saliva 2 h post-HM showed that six factors with 1 ≤ peptides and *p* < 0.05, were suppressed post-HM compared with their pre-HM levels, and the remaining 56 factors were unchanged (Table 2, Table S1). The candidate protein, CST4, is an enzyme of cysteine proteases that plays a protective function and regulatory role in the oral cavity (Dickinson D. P. et al., 2002). Salivary CST4 levels have been reported to specifically increase during ovulation in humans (Saibaba et al., 2016). Total cystatin activity in saliva is increased in patients with periodontal disease (Henskens et al., 1994), and CST4 possesses an antibacterial effect on *Porphyromonas gingivalis* involved in periodontal disease (Blankenvoorde et al., 1996). IGK, which is comprised of immunoglobulin, is the light chain (LC) and can be either a kappa or lambda isotype. LCs are involved in antigen recognition and activation of neutrophils and neurons (Brebner & Stockley, 2013). LCs can also be used as a marker for cancer and various diseases (Brebner & Stockley, 2013), and its serum expression levels have been reported to be suppressed in rheumatism, heart failure, diabetes, renal disease, intestinal disease, and HIV infection (Brebner & Stockley, 2013). According to recent reports, IGK and IG*λ* in saliva are suppressed after short-term intense training in highly trained male cyclists (Heaney et al., 2018). The suppression of IGK, but not IGλ, is in agreement with earlier studies by Heaney et al. (2018).

There are two likely causes of the differences between the study groups and exercise levels. Suppression of IGHA and IGK, components of IgA, may be involved in IgA function, and changes in specific LCs may imply regulation of inflammatory responses. Although the changes in IgA secretion and its involvement in the immune suppression state have been discussed (Campbell & Turner, 2018), to clarify the local immunity in the oral cavity and saliva in more detail, it would be helpful to examine not only the total amount of IgA but also its composition and activity.

KLK1 is a serine protease that is a component of the kallikrein kinin system (KKS) (Yiu et al., 2014). KLK1 cleaves low-molecular-weight kininogens into kinins, which perform biological functions via the kinin receptor signaling (Kakoki et al., 2009). KKS is involved in inflammation, coagulation, pain, and vascular permeability via kinin production. Kinin regulates cholinergic and sensory nerve stimulation by modulating bronchoconstriction, mucociliary secretion, and vascular permeability in the airway (Polosa R. et al., 1994). According to Polosa et al., asthmatics experience bronchoconstriction-induced bradykinin inhalation, but healthy subjects do not (Polosa R. et al., 1994). The effects of bradykinin in healthy subjects under exercise stress are unknown, but it is hypothesized that if KLK1 changes bronchioles as well as in the oral cavity, suppression of KLK1 may be related to bronchoconstriction during excessive exercise and may be involved in respiration.

In the oral cavity, KLK1 is secreted from the salivary glands, and *KLK1* expression in the parotid glands has also been reported (Saitou et al. 2020). Salivary KLK1 degrades neisserial heparin-binding antigen, a lipoprotein that is exposed on the surface of the meningococcal cell membrane (Pantano et al. 2019). To date, KLK1 behavior and mechanism during exercise stress has not been reported. Our study showed that KLK1 is suppressed under HM stress. For the reasons mentioned above, relative changes and the protein functions of CST4, KLK1, IGK and IGHA1, it is suggested that HM suppresses antimicrobial activity in the oral cavity.

IGHA1 has been used as a physical and mental stress marker (Pedersen et al., 2000). Moderate exercise increases sIgA, as does the activity of immune factors in the blood, and it decreases when the immune system is suppressed by HIE, such as in marathons (Nieman and Wentz, 2019). In terms of mental status, continued anxious mental states or transient nervousness decreases sIgA levels. On the other hand, a happy feeling and positive thoughts have been reported to increase sIgA levels (Obayashi K., 2013). Our results showed that CST4, KLK1, IGK, and IgA levels were similarly suppressed after the HM. A recent study reported that CST4 is activated and IGHA1 is suppressed during acute social stress in healthy young adults (Zallocco L. et al., 2021). According to recent reports, salivary proteins from highly trained athletes before and after competition increased the secretion of CST4 as a subacute response to aerobic and anaerobic exercises (Sant’Anna et al., 2019). Salivary free LC exhibits diurnal variation and is correlated with IgA (Rapson et al., 2020). Heaney et al. (2018) reported that salivary IGK and IGλ levels were suppressed after short-term intense training in highly trained male cyclists. As far as we have been able to determine, there had been no additional reports of IGK being changed by exercise. Similarly, there have been no reports of exercise or mental stress being associated with KLK1. The results of this and previous studies suggests that CST4 is activated during mental stress and suppressed during physical fatigue. Although sIgA has been suppressed during both mental stress and physical stress, CST4 may respond differently to mental stress and physical stress. Analysis of IGHA1 and CST4 behavior may provide a means of understanding mental and physical stress conditions in unknown saliva samples.

In this study, IGHA1, CST4, IGK, and KLK1 were detected in saliva proteins and were regulated by HM stress. Data from the Human Protein Atlas (www.proteinatlas.org) showed that the protein expression of IGHA1 has been reported in the intestine, lymphoid tissue, salivary glands, and stomach. Similarly, CST4 was reported in the salivary gland, IGK in the intestine and lymphoid tissue, and KLK1 in the pancreas and salivary gland.

We performed correlation analysis to determine whether there was a relationship between the regulated proteins (Fig. 5). Positive correlations were observed 2 and 4 h after HM for IGHA1, IGK, and CST4. These results suggest that protein regulation continued at 2 and 4 h. On the other hand, there was no correlation between IGHA1 and other proteins, suggesting that IgA and other proteins might be regulated through different pathways by HM stress. sIgA has been used as a marker of immunosuppression in exercise, but depending on the stress conditions of exercise, the same pathways and signals cannot be regulated by salivary proteins in the short period post-exercise. KLK1 after 2 h and IGK after 2 and 4 h were positively correlated (Fig. 5). The positive correlation between KLK1 and IGK may be difficult to discuss because our study did not observe protein localization or expression during HM stress.

However, these findings cannot be extrapolated to all groups of people. The limitations of this study are: (1) the study was conducted on women of a specific age group; (2) the study did not include men of the same age group; (3) this study did not assess exercise stress in detail. Although this study was conducted for the exercise stress of completing an HM, the information obtained in the experiment was not sufficient to analyze the exercise stress state in detail, and the detailed exercise stress state of each individual was unclear. Further studies are required to address these issues.

## Conclusions

HM reduced saliva flow rate and IGHA1 secretion. The proteome and WB analysis showed that CST4 and KLK1 suppressed 2 h post and 4 h post-HM, IGK suppressed in 2 h post-HM. In addition, IGHA1, CST4 and IGK protein were positive correlation at 2 h and 4 h post-HM, suggested that protein regulation remains at 2 h and 4 h post-HM as same. Our findings of this study can be applied to health management of recreation runner and people who engage in high to middle intensity physical activities in daily life.

## Acknowledgements

We would like to acknowledge Dr. Kazuhiro Sako for critiques of our experiment. We would like to thank of Yuki Miura and Kumi Hasegawa who assisted with implementation of this experiment.

## Figure Legends

**Supplemental Fig. 1 SDS-PAGE analysis of salivary proteins.**

A total of 3 μg protein was loaded from each sample. The protein samples in the NExG were from the first, second, and third sampling, while those in the ExG were from pre-HM and 2 h post and 4 h post-HM. Protein samples from a single subject were used in NExG and ExG. The protein bands with asterisks in the ExG decreased 2 h post and 4 h post-HM compared with pre-HM.

**Supplemental Fig. 2. Venn diagram of the number of quantified proteins and Protein distribution of molecular weight and fold change.**

(A) Venn diagram of the number of quantified proteins derived from the five sets of iTRAQ experiments. (B) Protein distribution in accordance with log_10_ theoretical molecular weight versus log_2_ fold change of all the detected proteins of 1 ≤ peptide.

**Supplemental table 1. List of salivary proteins identified by iTRAQ analysis in pre- and post-half marathon.**

The peptide shows the maximum detected number in analyses samples. *The subcellular location and tissue specificity shows the localization of detected proteins. Data were from the human protein atlas (https://www.proteinatlas.org/). n.d; no data. **The mRNA was described as to specifically expressed in any of the salivary glands. The sites of adult salivary glands that were more than twice as active as fetal salivary glands with P<0.05 were shown. ms; submandibular gland, PAR; parotid gland, SL; sublingual gland. The data of mRNA localization were from Saitou et al 2020. ***The whole saliva indicates whether the detected protein is contained in the salivary proteins of the Human Salivary Protein Wiki (https://salivaryproteome.nidcr.nih.gov/). Y; contain in saliva, -; no existence.

