## Supplementary figures and images for "Analysis of IGHA1 and other salivary proteins post half marathon in female participants"

### Supplemental fig

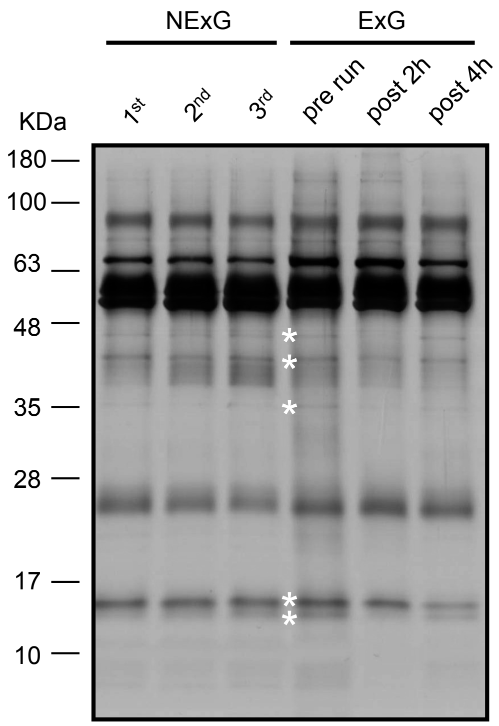


Maruyama et al sFig.1

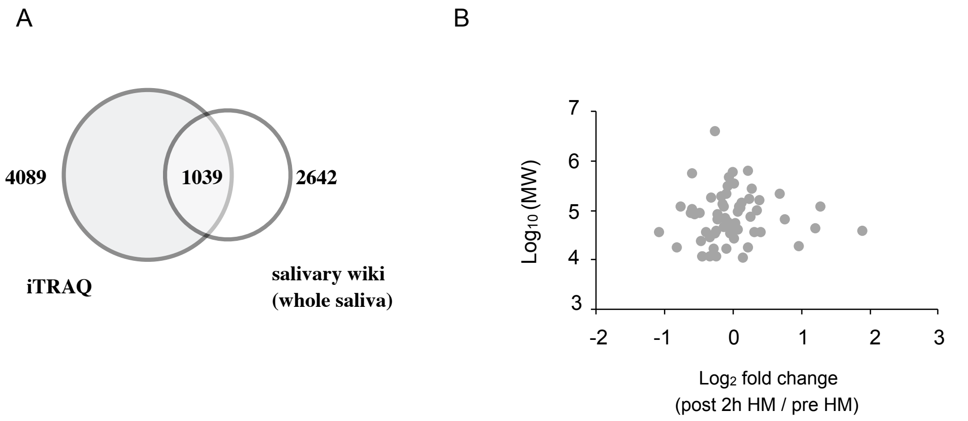


Maruyama et al sFig.2
